# DNAJA2 and Hero11 mediate similar conformational extension and aggregation suppression of TDP-43

**DOI:** 10.1101/2022.11.15.516569

**Authors:** Andy Y.W. Lam, Kotaro Tsuboyama, Hisashi Tadakuma, Yukihide Tomari

**Author notes:** To whom correspondence should be addressed.; Correspondence may also be addressed to Kotaro Tsuboyama.; or Hisashi Tadakuma.

## Abstract

Protein misfolding and aggregation are characteristic features of neurodegenerative diseases. While molecular chaperones are well-known suppressors of these aberrant events, we recently reported that highly disordered, hydrophilic and charged heat-resistant obscure (Hero) proteins may have similar effects. Specifically, Hero proteins can maintain the activity of other proteins from denaturing conditions in vitro, while their overexpression can suppress cellular aggregation and toxicity associated with aggregation-prone proteins. However, it is unclear how these protective effects are achieved. Here, we utilized single-molecule FRET to monitor the conformations of the aggregation-prone prion-like low complexity domain (LCD) of TAR DNA-binding protein 43 (TDP-43). While we observed high conformational heterogeneity in wild-type LCD, the ALS-associated mutation A315T promoted collapsed conformations. In contrast, an Hsp40 chaperone, DNAJA2, and a Hero protein, Hero11 stabilized extended states of the LCD, consistent with their ability to suppress the aggregation of TDP-43. Our results link single-molecule effects on conformation to macro effects on bulk aggregation, where a Hero protein, like a chaperone, can maintain the conformational integrity of a client protein to prevent its aggregation.

## Introduction

Most protein functions are defined by their native state, usually consisting of well-folded tertiary structures. However, intrinsically disordered proteins (IDPs) do not follow this principle. Particularly when isolated, IDPs exhibit a high degree of conformational flexibility, adopting a range of loosely defined structures owing to the high hydrophilicity of their amino acid compositions and the absence of a hydrophobic ’core’ necessary for stabilizing tertiary structures. This flexibility enables cooperative, multivalent interactions with various binding partners in different contexts. Consequently, IDPs play diverse biological roles, including gene regulation, signal transduction, trafficking, and assembly (1, 2).

Despite these important functions, several IDPs have become the focus of intense study because of their aberrant behavior in neurodegenerative diseases. One representative example of such a misbehaving IDP is transactive response DNA binding protein 43 (TDP-43), a highly abundant and essential RNA-binding protein that functions in various manners of RNA processing, including mRNA transcription, splicing, transport and translation, and the processing of microRNAs and long noncoding RNAs (3–5). TDP-43 is predominantly localized to the nucleus but can move into the cytoplasm upon cellular stress, including oxidative, osmotic, heat, proteasome, and ER stresses, where it not only regulates but also forms stress granules (6, 7). During prolonged stress, TDP-43 even forms aberrant aggregates in the cytosol (8–11). These aggregates are known as pathological hallmarks of amyotrophic lateral sclerosis (ALS) and appear in ∼97% of cases (12, 13) and in ∼43% of cases of frontotemporal lobar degeneration (FTLD) (3). In these cases, the ∼141 amino acid intrinsically disordered prion-like C-terminal region of TDP-43, called the low-complexity domain (LCD), forms the tightly packed stabilizing cores of these aggregates. (14–17).

To counteract the disasters caused by protein aggregates, cells have developed mechanisms to suppress misfolding; for example, molecular chaperones that mainly assist in the proper folding of proteins during or after translation (18). Chaperones usually work in an ATP-dependent manner, promoting the proper folding of unfolded client proteins or the disassembly of already aggregated proteins, for example, the Hsp70/40/90 and the Hsp70/40/110 systems, respectively (18–20). Additionally, chaperones can also work in an ATP-independent manner by binding to unfolded client proteins to prevent their aggregation. An example of this are Hsp40s, also known as DnaJ or J-domain proteins, which bind and stabilize unfolded client proteins until they are handed off to Hsp70 (20–22). In both ATP-dependent and ATP-independent cases, chaperones bind the folding intermediates of their client proteins through mainly hydrophobic interfaces on the tertiary structure of the chaperones (23–25). Chaperones collaborate with each other and other processes, such as degradation pathways, to form an integral part of the protein quality control network (18, 20).

In contrast to structure-based conventional chaperones, studies of late-embryogenesis abundant (LEA) proteins in plants, cytoplasmic-abundant heat-soluble (CAHS) proteins in tardigrades, and other hydrophilic IDPs suggest that they may also have chaperone-like activities (2, 26–35). Additionally, we previously reported that heat-resistant obscure (Hero) proteins, which are abundant in the boiled supernatant of human and fly cell lysates, can stabilize the activity of client proteins from denaturing conditions in vitro and prevent pathological protein aggregation in cells, suggestive of their potential in biotechnology and therapeutics (36, 37). Moreover, overexpression of some Hero proteins can extend the life span of *Drosophila* by >30% (36). However, hero proteins appear vastly different from structured chaperones in that they are highly hydrophilic, electrostatically charged, and intrinsically disordered. Remarkably, Hero protein activity seems to rely on their biophysical properties as highly charged and sufficiently long polymers, rather than specific sequence-encoded structure (36). In light of these obvious structural differences and unique biophysical properties compared to canonical chaperones, it is important to investigate the mechanism of how these IDPs suppress aggregation.

Here, we report unexpectedly similar effects of a hero protein, Hero11, and an Hsp40 family chaperone, DNAJA2, on the conformational dynamics and aggregation of TDP-43. Using genetic code expansion in HEK293T cells, we synthesized dye-labeled full-length TDP-43 for monitoring its LCD via total internal reflection fluorescence (TIRF) single-molecule fluorescence energy transfer (smFRET). While we observed high heterogeneity in the conformation of the WT LCD, the disease-associated aggregation-enhancing A315T mutation promoted collapsed conformations. In contrast, both DNAJA2 and Hero11 stabilized extended conformations, while the chargeless Hero11 mutant did not, mirroring their effects on aggregation. This suggests that Hero11, despite lacking clear structural features like conventional Hsp40, can still stabilize extended conformations resulting in a similar effect.

## Materials and Methods

### HEK293T cell culture and lysate preparation

HEK293T cells were maintained in Dulbecco’s Modified Eagle Medium (DMEM; Sigma, QJ-D6429) supplemented with 10% FBS in a 5% CO_2_ atmosphere at 37°C. Naïve HEK293T cell lysates were produced as previously described (36). Briefly, HEK293T cells were grown to confluency in 10 ∼ 20 100 mm plates (Corning), harvested and resuspended in 1 μL 1× hypotonic lysis buffer (10 mM HEPES KOH, 10 mM KOAc 1.5 mM Mg(oAc)C containing 1× cOmplete™ EDTA-free Protease Inhibitor Cocktail (Roche) and 1 mM DTT on ice to lyse cells, and lysate collected as the supernatant from 20 min 17, 000 g centrifugation at 4°C.

### Plasmids for recombinant protein production in E. coli

Plasmids for recombinant Hero proteins and the controls, pColdI_ Hero11-FLAG-6×His, Hero11KR→G-FLAG-6×His, and GST-FLAG-6×His, were previously described (36). The recombinant plasmid used for the chaperone pColdI_6×His-DNAJA2 was previously described (38).

### Plasmids for HEK293T cell-based assays and protein production

For transfection into HEK293T cells, pCAGEN_6×His-DNAJA2-HA was generated via PCR of the ORF from pColdI_6×His-DNAJA2 and inserted into EcoRI-digested pCAGEN plasmids using NEBuilder HiFi DNA Assembly Master Mix (NEB). This plasmid was subsequently used to produce pCAGEN_3xFLAG-DNAJA2 in a similar manner. The plasmids used for the Hero protein and the controls pCAGEN_3xFLAG-Hero11, pCAGEN_3xFLAG-Hero11KR→G, and pCAGEN_3xFLAG-GST were previously described (36). The plasmids used for the chaperones pCAGEN_3xFLAG-DNAJB8 and pCAGEN_3xFLAG-DNAJC8 were generated as previously described (39). The plasmids used for genetic code expansion, pCAGEN_U6tRNAUCmut×4_NESPylRS-Afmut and pCAGEN_U6tRNAUCmut×4_eRF1EDmut, were previously described (40). The plasmid used for the expression of GFP-tagged TDP-43 in HEK293T cells and TDP43ΔNLS was previously described (36). The plasmid for incorporating unnatural amino acids into TDP-43, pCAGEN_3xFLAG-TEV-HALO-TDP43_R268Amb-F420Amb-2×HA, was created first by PCR of the ORF from pCAGEN_eGFP-TDP43 and insertion into NotI-digested pCAGEN_FLAG-tev-HALO-Ago2 (40). Then, an ochre mutation of the final stop codon after 2×HA, and amber mutations around the TDP-43 LCD at R268 and F420 were introduced by PCR-based site-directed mutagenesis, as previously described. To test the A315T mutation, pCAGEN_eGFP-TDP43_A315TΔNLS and pCAGEN_FLAG-TEV-HALO-TDP43_R268Amb-A315T-F420Amb-2×HA were created from the A315T mutation of pCAGEN_eGFP-TDP43_ΔNLS and pCAGEN_FLAG-TEV-HALO-TDP43_R268Amb-F420Amb-2×HA, respectively, via PCR-based site-directed mutagenesis.

### Protein purification of His-tagged proteins from pColdI plasmids

Recombinant Hero proteins, charge mutants and DNAJA2 were purified as previously described (36, 38). Briefly, pColdI plasmids for C-terminally FLAG-6×His-tagged proteins were transfected into Rosetta2 (DE3) cells, which were subsequently grown at 37°C to an O.D. of 0.4−0.6. The proteins were induced with 1 mM isopropyl-β-D-thiogalactoside (IPTG) overnight at 15°C and snap-frozen with liquid nitrogen. Cells were lysed by probe sonication in His A buffer (30 mM HEPES-KOH pH 7.4, 200 mM KOAc, 2 mM Mg(OAc)2, 5% glycerol) supplemented with cOmplete™ EDTA-free Protease Inhibitor Cocktail (Roche), cleared by centrifugation, purified over Ni-NTA resin (Roche) and eluted with His B buffer (HisA with 400 mM imidazole). The proteins were buffer-exchanged into 1×PBS using PD-10 columns (Cytiva), and the concentration was determined by a BCA assay (Pierce) before supplementation to 1 mM DTT. The proteins were snap frozen in aliquots and stored at −80°C.

### Protein purification via genetic code expansion

Genetic code expansion was performed as described previously (40–42). HEK293T cells were seeded and transfected after 24 h with pCAGEN plasmids in medium supplemented with EndoBCN-K (SiChem, SC-8014) and harvested 48 h after transfection. The cells were lysed by sonication via BIORUPTOR II (BM Bio), snap-frozen with liquid nitrogen and stored at −80°C. The proteins were purified with FLAG-functionalized Dynabeads Protein G (Invitrogen), dye-labeled on beads with tetrazine-functionalized Cy3 and ATTO647N (Jena Bioscience, CLK-014-05 and CLK-012-02), functionalized with the HaloTag Biotin ligand (Promega, G8281), eluted in naïve HEK293T cell lysate with TurboTEV Protease (Accelagen, T0102S), and snap-frozen in liquid nitrogen and stored at −80°C. Fluorescent dye labeling was verified via SDSCPAGE via detection of fluorescence at the indicated wavelengths with an Imager 600 (Amersham).

### Cell-based aggregation assays

Cell-based aggregation assays were performed as previously described (36). HEK293T cells were seeded in 12-well plates (Corning) at a density of 6×10C cells/mL, and after 24 h co-transfected with 200 ng of the pCAGEN plasmid for TDP-43 and 1000 ng of the pCAGEN plasmid for the other protein using Lipofectamine 3000 Transfection Reagent (Invitrogen, L3000001). After 48 h, cells were imaged using a FLUOVIEW FV3000 confocal laser scanning microscope (Evident) and then lysed with BIORUPTOR II (BM Bio) in 1×PBS supplemented with 1× cOmplete™ Protease Inhibitor Cocktail (Roche) in Protein LoBind Tubes (Eppendorf). Total protein amounts were quantified via Bradford assay (Bio-Rad) normalized to the lowest concentration sample. Samples were solubilized in equal volume of 1×PBS with 1% SDS and run through a 0.2 μm Whatman OE66 cellulose acetate membrane filter (GE Healthcare, 10404180) using a suction pump (Air Liquide) at a vacuum of around 10 kPa, to capture aggregated proteins. TDP-43 aggregation was checked using GFP fluorescence signal, and for the final figures, detected via immunoblotting for GFP using an α-GFP (B-2) primary antibody (Santa Cruz Biotechnology, sc-9996) and an α-mouse IgG secondary antibody (MBL), both at 1:10000.

### Microscopy of GFP-TDP43ΔNLS in cells

HEK293T cells were seeded at a density of 6×10C cells/mL onto 35 mm glass bottom dishes (MatTek, P35G-1.5-14-C) pretreated with poly-L-lysine (SigmaCAldrich, P4707), and after 24 h were transfected with 1000 ng of pCAGEN_GFP-TDP43ΔNLS and 1500 ng of pCAGEN-3xFLAG-DNAJA2, Hero11, Hero11KR→G, or GST using Lipofectamine™ 3000 Transfection Reagent (Invitrogen, L3000001). After 48h, cells were stained with Hoechst 33342 (Invitrogen, H3570) for imaging on a FLUOVIEW FV3000 confocal laser scanning microscope (Evident) using a 100x oil-immersion lens controlled with FV31S-SW Viewer software (Evident). All images were captured using the same laser power and settings, chosen as to avoid saturation of pixels. For each image, reference images with saturation of pixels were also captured.

For FRAP experiments, cells were transfected with 1000 ng of pCAGEN_GFP-TDP43ΔNLS or 1500 ng of pCAGEN-3xFLAG-GST without Hoechst staining using the LSM stimulation mode. Zoomed-in videos of a single cell were taken at 1 second per frame. After five pre-bleaching frames, 100% power laser stimulation was applied to a small circular area within the puncta for 500 ms, and an image was taken as the bleach frame, followed by 150 additional recovery frames.

### Single-molecule FRET microscopy

Single-molecule assays were conducted as previously described (40, 43), with modifications to minimize aggregation. Briefly, aliquots of dye-labeled HALO-TDP43 were prediluted in 1× lysis buffer (30 mM HEPES-KOH pH 7.4, 100 mM KOAc, 2 mM Mg(OAc)_2_) and then diluted in naïve HEK293T cell lysate with 1×ATP regeneration system (1 mM ATP, 25 mM creatine monophosphate (Sigma), 0.03 U/μL creatine kinase (Calbiochem), 0.1 U/μL RNasin Plus RNase Inhibitor (Promega), in lysis buffer) and incubated at 37°C for 30 min to enable aggregation-clearing chaperone activities. Note that the use of high-quality HEK293T cell lysates generated via the hypotonic lysis method was crucial, as lysates prepared via sonication failed to sufficiently remove aggregates. The mixture was subsequently centrifuged at 18, 000×g for 15 min at 25°C, after which the supernatant was collected to remove the aggregated proteins in the pellet. Observation chambers were prepared as previously described from biotin-PEG- and PEGylated glass slides (40, 43), infused with neutravidin for 2 min, and washed with 1×lysis buffer. The supernatant was added to the chamber, which was incubated for 2 min, washed twice with wash buffer (1× lysis buffer containing 1% Triton X-100 and 800 mM NaCl), four times with wash buffer supplemented with 10 mM ATP, and four times with 1×lysis buffer. These stringent washes are necessary to remove nonspecific interactions and oligomerization of TDP-43 with tethered TDP-43 molecules. The final observation buffer containing 0.3 mg/mL BSA or recombinant protein in 1×PBS with 1 mM DTT, an O_2_ scavenger system (7.1 mM PCA (3, 4-dihydroxybenzoic acid, 37580, Sigma), 29 nM PCD (protocatechuate 3, 4-dioxygenase, P8279, Sigma) (44), and 2 mM Trolox (6-hydroxy-2, 5, 7, 8-tetramethylchroman-2-carboxylic acid, 238813, Sigma) was flowed into chamber, after which the chamber was mounted onto a custom microscope for smFRET observation. For the experiments at 0.075 mg/mL, 0.075 mg/mL of the recombinant protein was supplemented with BSA for a final total protein concentration of 0.3 mg/mL. Three five-minute videos were taken of each slide using alternating laser illumination of 1 s each. We emphasize that these procedures are uniquely practical for TIRF-smFRET experiments, which, by tethering dye-labeled molecules, allow for buffer exchange and minimization of aggregation for an extremely small sample.

### Data analysis and visualization software

Image analyses of cells were performed using the Fiji distribution of ImageJ (45). The data analysis was performed in Python 3.7.10 using compatible versions of pandas, NumPy, SciPy, Matplotlib, and seaborn in addition to the standard library via Jupyter notebooks implemented with Spyder-notebook in Spyder IDE version 5.

### Analysis of aggregation assays

Images from bulk aggregation filter trap assays were taken using an Amersham ImageQuant 800 Imager (Amersham) in TIFF image format. For quantification, images were analyzed using ImageJ Fiji using a simple macro script in a manner similar to dot blots, first with background subtraction via the “Subtract Background” tool, with a rolling ball radius of 50 pixels. Circular areas of 125 pixels diameter were then selected around the highest concentration wells to obtain integrated densities for each sample and saved in CSV file format. These measurements were imported into Python for analysis, where integrated densities were divided by the GST co-transfection well in each blot to quantify aggregation relative to GST and visualized.

### Microscope image analysis

The HEK293T cell images in the figures were prepared via the QuickFigs plugin for Fiji (46). For puncta counting and measurement analyses, images were analyzed through an automated ImageJ macro that used base macro functions to threshold, clean, and identify areas of interest. These created masks for cell areas, nuclei areas, and puncta areas, and image calculator operations were performed to identify overlapping areas. Cell area masks were defined using the saturated images to identify cells with any expression of GFP, including diffuse background signals. Nuclei areas masks were defined using the using nuclei that were contained in the cell area mask and were also used as an accurate count of cells of interest. Puncta area masks were defined using masks that detected local areas of high and moderate GFP signals and thresholding against the low background signal. Using these masks cytoplasmic puncta were defined in order to count and measure areas and signals of these puncta in the original image and exported to CSV for analysis and visualization in Python.

### FRAP analysis of large aggregates

For FRAP analysis, an interactive macro was used to for allow partially automatic selection of regions of interest via local maxima and thresholding: the bleach spot, the puncta area, the cell area and a background area, and were used for FRAP analysis. These regions were interpolated over the video to account for movement of the puncta and cells and were subsequently measured to determine mean signal in the area and exported to csv for analysis in Python. The measurements for each recovery curve were background corrected and double normalized to the average maximum intensity pre-bleach and minimum intensity post-bleach (47), and visualized. A single exponential, exponential equation (Equation 1) was fitted to each recovery curve using non-linear least squares via SciPy for determination of the mobile fraction (1-I_f_).

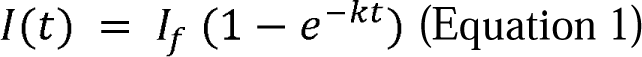

where the intensity at time *t, I(t)*, is described by the rate parameter k and I_f_ is the plateau recovery intensity.

### Analysis of smFRET assays

The smFRET videos were imported to iSMS implemented in MATLAB for image channel alignment, peak localization, FRET-pair identification, photobleaching and blinking identification, drift correction, and correction factor determination and application as previously described (40, 48, 49). Note that background correction, applied uniformly by experiment, sometimes results in negative FRET values close to 0. FRET efficiency traces were selected based on single step photobleaching, a stoichiometry of ∼0.4 via ALEX and anticorrelation. The FRET distribution analysis data were exported as TXT files from iSMS. Visualization was conducted through python to load TXT files, and density histograms were plotted using the combined data for each condition with 25 bins. Average density plots were created by estimating the probability density functions for each independent experiment via Gaussian kernel density estimation with 0.05 bandwidth, and the values of each curve were evaluated for determination of means and standard errors along the entire range. The mean density and standard error of the mean for each condition were plotted over the histograms.

### Analysis of smFRET dynamics

For dynamics analysis, a hidden Markov model (HMM) with variational Bayesian expectation maximization (VBEM) implemented in iSMS was used to extract ideal traces, as previously described (40, 49), as TXT and imported into python for analysis. Traces were denoised for counting transition events and dwell times and visualized. Single exponential distributions were fitted to the dwell time distributions using Bayesian modeling implemented in PyMC 5.9. A normally distributed (μ=0, σ=10) prior was used for the log of the lambda parameter (A, equivalent to the rate parameter, k) for the exponential model and combined with the observed dwell time distribution in the likelihood function to run Markov chain Monte Carlo (MCMC) stochastic simulations with the No U-Turn Sampler (NUTS) algorithm, taking 100 samples after 100 tuning samples from one chain. We reported median and 95% credible interval estimates of the posterior distribution of the fitted parameters. These were visualized as cumulative distribution functions (Equation 2).

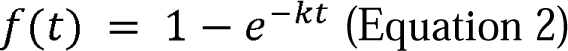

where the cumulative distribution function f(t) is described by the rate parameter k.

### Inter-dye distance estimations from FRET efficiencies

To approximate theoretical end-to-end distances from FRET efficiencies, we used community-provided values of the Förster radius R_O_ for FRET between Cy3 and ATTO 647 N (50) following Förster’s equation for the relationship between FRET efficiency and distance.

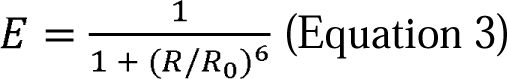

where the FRET efficiency, E, is proportional to the 6^th^ power of the distance between fluorophores, R.

### Wormlike chain model

To approximate the end-to-end distance, we assumed that the LCD polymer behaves according to the wormlike chain (WLC) model (51–53).

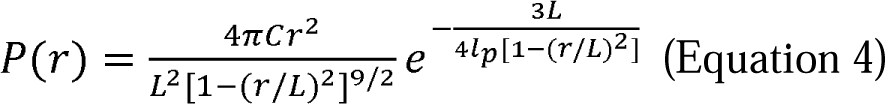

where r is the distance between the polymer ends, L is the polymer contour length, P(r) is the probability of the polymer having an end-to-end distance of r, l_p_ is the persistence length of the polymer (describing bending), and C is the normalization factor. For simplicity, we considered a one-dimensional distribution with a contour length of 47.94 nm (141 amino acids × 0.34 nm per amino acid). A persistence length of 0.8 nm was chosen to describe random coils, as experimentally observed for poly-glycine (or glycine– serine repeats) (51).

## Results

### DNAJA2 and Hero11 suppress TDP-43 aggregation in HEK293T cells

To understand how Hero proteins act upon aggregation, we first sought to identify a well-established chaperone protein for reference. Since we do not expect Hero proteins to use ATP for this activity, we reasoned that the ATP-independent chaperoning ability of Hsp40 proteins would be a suitable reference for studying Hero proteins. Thus, we first performed cell-based aggregation assays to identify an effective Hsp40 chaperone for the aggregation of TDP-43 (36). As in the previous study, we used GFP-tagged TDP-43 lacking its nuclear localization signal (GFP-TDP43ΔNLS WT) to increase cytoplasmic mislocalization and aggregation (54, 55). GFP-TDP43ΔNLS was co-transfected with Hero proteins, Hsp40 proteins, or GST (Glutathione S-transferase) control into HEK293T cells, and aggregation was assessed by the filter trap method (Figure 1A). In agreement with our previous results, for an 11 kDa highly positively charged Hero protein, Hero11 (also known as the L10K protein or C19orf53), strongly suppressed TDP-43 aggregation, while the KR→G charge mutant, in which all 21 lysine and 6 arginine residues were mutated to neutral glycine, failed to prevent aggregation (Figure 1B−D), suggesting the importance of the net charge for this anti-aggregation activity, as shown previously (36). Of the tested Hsp40s, DNAJB8 and DNAJC8 did not appear to reproducibly suppress aggregation in the three experimental replicates, and notably DNAJB8 suppresses aggregation signals in microscope images but not in the filter trap assay. This reduction of GFP signal in microscope images may be attributable to denaturation of GFP or quenching of the fluorophore which does not affect the size- and antibody-based filter trap assay. However, DNAJA2 most effectively and reproducibly suppressed the aggregation of co-transfected GFP-TDP43ΔNLS relative to the GST control in both microscope and filter trap assay (Figure 1B−D). Given that DNAJA2 was recently demonstrated to selectively bind to and reduce the phase separation of TDP-43 LCDs (56), our results support DNAJA2 as a representative suppressor of TDP-43 aggregation. Thus, we chose DNAJA2 for further analysis alongside Hero11.

**Figure 1.**
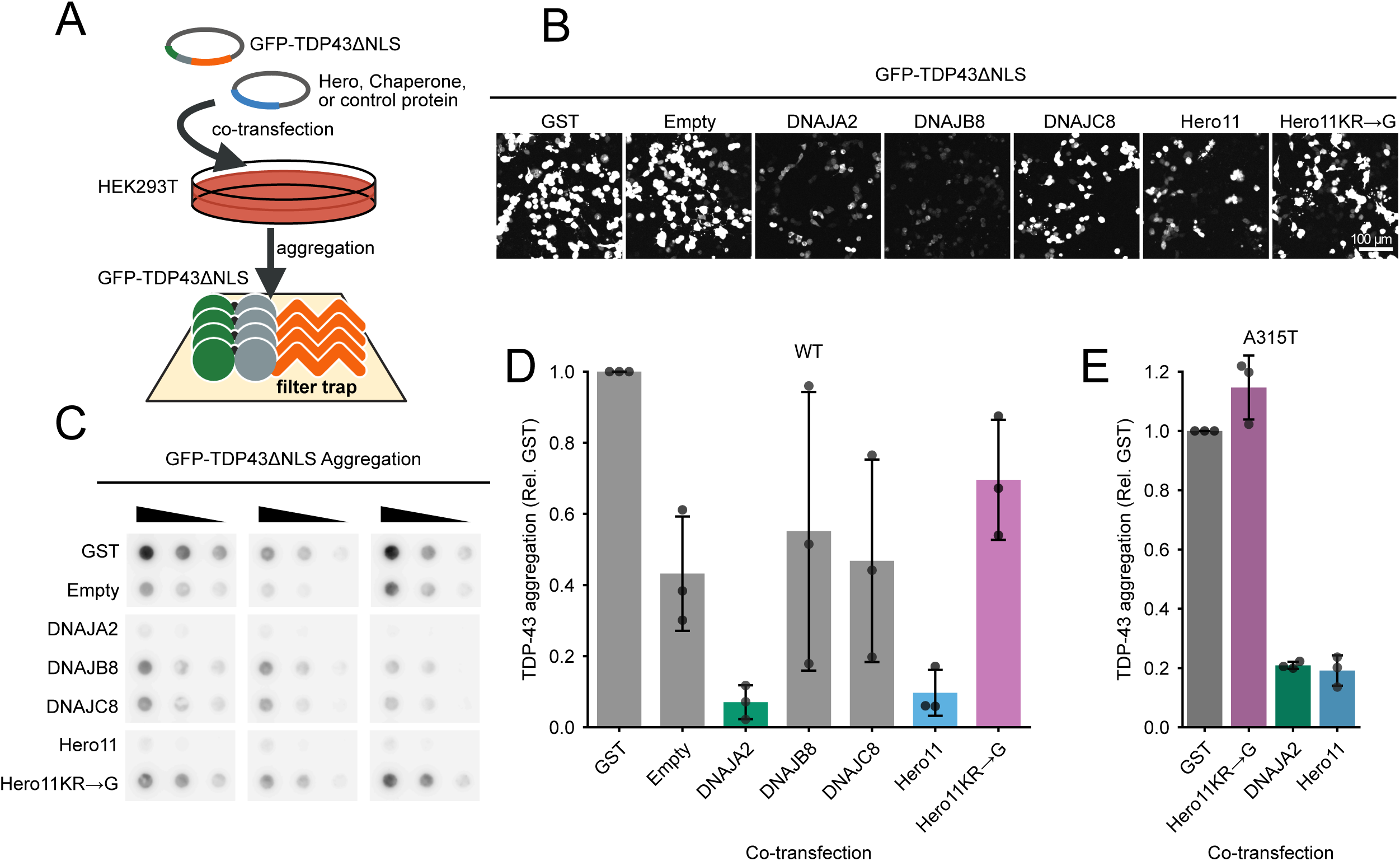
DNAJA2 and Hero11 suppress the aggregation of TDP-43 in HEK293T cells. (A) Schematic of the HEK293T cell-based filter trap assay for TDP-43 aggregation. Plasmids for GFP-TDP43ΔNLS and the protein indicated in (B) were co-transfected into HEK293T cells and incubated for 48 h. The cells were subsequently lysed and filtered through a 0.2 μm filter membrane under vacuum, on which aggregated proteins become trapped for detection via immunoblotting for GFP. (B) Representative microscope images of GFP-TDP43ΔNLS signals in HEK293T cells co-transfected with the indicated chaperones, hero proteins or control at 20x optical magnification. Empty refers to the pCAGEN plasmid without any insert protein. (C) Immunoblots for aggregated GFP-TDP43ΔNLS in HEK293T cells detected with α-GFP antibody. For each filter trap membrane (from three independent experiments), each sample was loaded three times at two-fold serial dilution. (D) Quantification of GFP-TDP43ΔNLS WT aggregation density in (C). For each membrane, values are normalized to the aggregation in the GST co-transfection. Bar plots represent the mean and standard deviation as the bar height and error bar, respectively. Dots represent the values of individual experiments. (E) Quantification of GFP-TDP43ΔNLS A315T aggregation density normalized to aggregation in GST co-transfection for each membrane (from three independent experiments). Bar plots represent the mean and standard deviation as the bar height and error bars, respectively. Dots represent the values of individual experiments.

Many disease-associated mutations of TDP-43 have been found in the LCD, for example, A315T, which stabilizes amyloid fibrils (16, 17) and promotes its aggregation and toxicity (57). Thus, we next introduced the A315T mutant into GFP-TDP43ΔNLS (GFP-TDP43ΔNLS A315T) and co-transfected it with Hero11 or DNAJA2. Similar to the WT GFP-TDP43ΔNLS, DNAJA2 and Hero11, but not Hero11KR→G, suppressed the aggregation of A315T (Figure 1E), suggesting that both of these proteins are effective suppressors of the pathogenic A315T mutant.

We next performed detailed microscopy analysis to investigate how Hero and DNAJA2 affect the morphology and dynamics of GFP-TDP-43ΔNLS aggregates in HEK293T cells. While large, irregularly shaped puncta of GFP-TDP43ΔNLS appeared in the cytoplasm of cells co-transfected with GST or Hero11KR→G, these puncta were absent in the DNAJA2 and Hero11 co-transfections which instead contained diffuse cytoplasmic signals (Figure 2A). To quantify this difference, we used an automated macro script to identify puncta signals from microscope images, from which puncta counts and areas were measured. Our image analysis pipeline revealed that cells co-transfected with GST and Hero11KR→G contained ∼10 μm^2^ of total puncta signal area on average, but DNAJA2 and Hero11 co-transfections reduced this area to ∼0.3 and ∼0.8 μm^2,^ respectively (Figure 2B).

**Figure 2.**
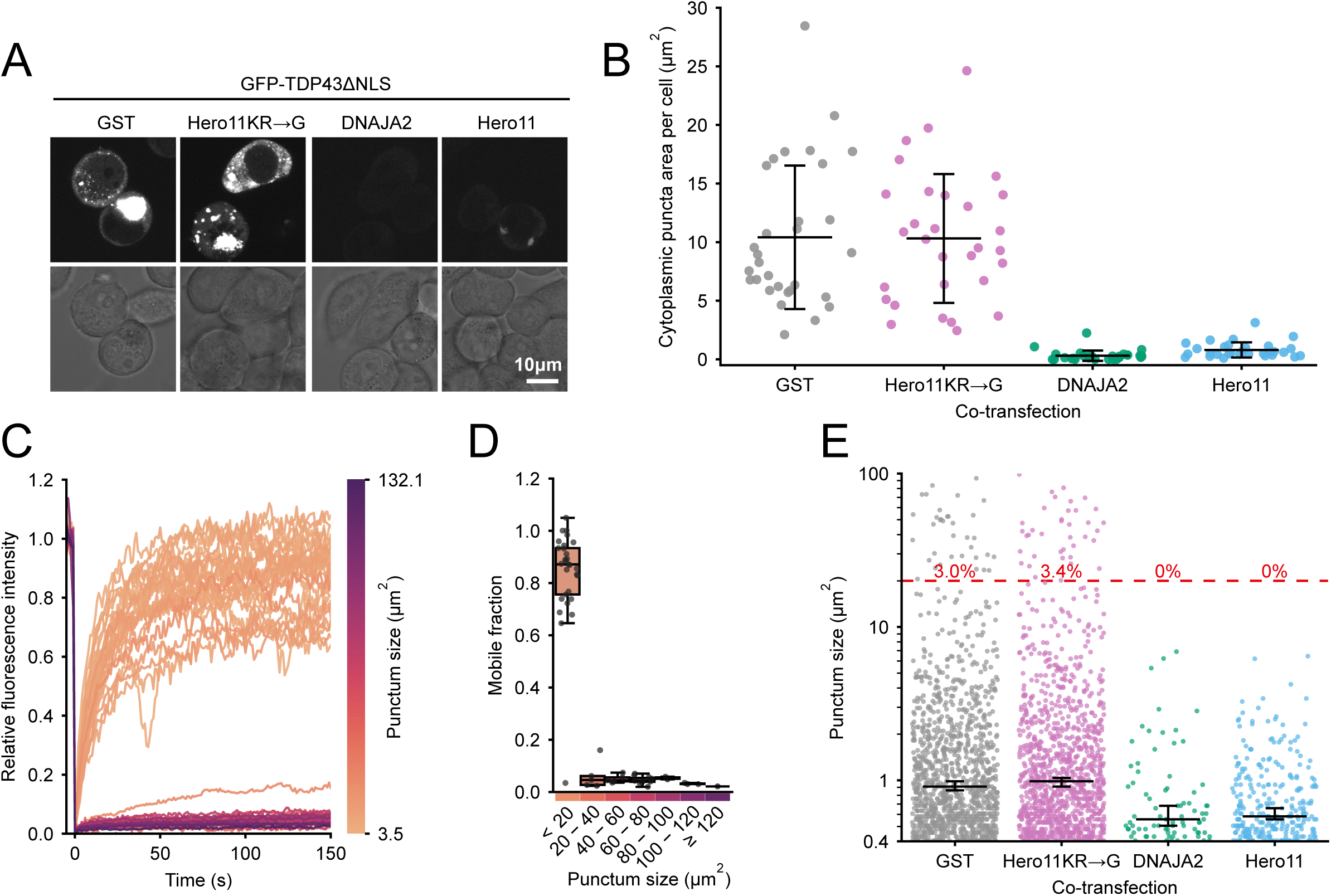
DNAJA2 and Hero11 suppress large immobile aggregates of TDP-43. (A) Representative microscope images of GFP-TDP43ΔNLS signals in HEK293T cells. Cells were co-transfected with plasmids for GFP-TDP43ΔNLS and the indicated proteins, and after 48h imaged at 100x optical magnification. Top row, GFP-TDP43ΔNLS signals. Bottom row, brightfield image. (B) Total area (µm²) of GFP-TDP43ΔNLS puncta in the cytoplasm per cell. For each image, total puncta area was measured and normalized by dividing over the number of cells. Each dot represents one image. The error bars represent the means and standard deviations calculated from the 30 images per co-transfection condition. (C) FRAP recovery curves of GFP-TDP43ΔNLS puncta. Cells were co-transfected with plasmids for GFP-TDP43ΔNLS and GST, and after 48h FRAP was performed via bleaching of an area in the middle of each punctum. Each trace represents the recovery curve for a punctum from −5 s pre-bleach to 150 s post-bleach (where the area within the punctum was bleached at 0 s) and colored according to puncta size. (D) Punctum internal mobility binned according to punctum size. The mobile fractions were extracted from the fits of the exponential recovery functions to the FRAP recovery curves for each punctum. Each dot represents a punctum. Boxplots represent the median and interquartile range, and whiskers represent the 1.5x interquartile range. (E) Cytoplasmic punctum sizes in each of the co-transfections. Each dot represents a punctum. Error lines represent the median and 95% confidence intervals, chosen due to the asymmetrical distribution, calculated from all puncta of a co-transfection. The dotted line indicates 20 µm², and the red text indicates the percentage of aggregates in each co-transfection above this line. There are 1156, 1107, 79, and 281 puncta for the GST, Hero11KR→G, DNAJA2 and Hero11 co-transfections, respectively.

To understand the biophysical properties of these puncta, we performed fluorescence recovery after photobleaching (FRAP) of GFP-TDP-43ΔNLS bodies in the GST control. Under our experimental conditions, we observed a discontinuous relationship between the size of the puncta and internal mobility, where small puncta with areas <20 µm² tended to have recoveries >0.8, while larger bodies with areas ≥20 µm² had <0.1 recovery (Figure 2C and 2D). This suggests that while smaller puncta tend to be highly fluid, bodies larger than 20 µm² are practically immobile. To understand how DNAJA2 and Hero11 reduce these puncta, we also quantified the sizes of individual puncta in the images (Figure 2E). The sizes were unevenly distributed in all co-transfections, consisting of ∼50–75% small areas ≤1 µm², for all four conditions we tested here. However, in the GST and Hero11KR→G co-transfections, puncta tended to be larger and more numerous compared to those in DNAJA2 and Hero11. Notably, while ∼3% of the puncta in the GST (35 out of 1156) and Hero11KR→G (38 out of 1107) co-transfections were larger than 20 µm², we observed no puncta above this size threshold out of 79 and 281 total puncta in the DNAJA2 and Hero11 transfections, respectively (Figure 2E). Taken together, these findings suggest that both Hero11 and DNAJA2 efficiently suppress the transition of small liquid-like TDP-43 bodies into larger solid cytoplasmic aggregates, which may be toxic and cause disease (11, 58).

### The TDP-43 LCD takes on heterogenous and compact conformations

Given the importance of the intrinsically disordered LCD for TDP-43 aggregation (14–16), we next sought to observe the conformational behavior of this region at the single-molecule level. Total internal reflection fluorescence (TIRF)-based smFRET is a powerful tool for isolating and collecting long-range conformational data of individual IDPs that are difficult to obtain by other methods (59–61), resolving heterogeneity between molecules. Using genetic code expansion in HEK293T cells, we synthesized full-length TDP-43 with the unnatural amino acid endo-BCN-L-lysine (endoBCN-K) incorporated around the LCD via the mutation of R268 six amino acids before and in a linker six amino acids after the LCD (Supplementary Figure 1). We then performed site-specific dye labeling at these two positions with tetrazine-coupled fluorescent dyes Cy3 (donor) and ATTO647N (acceptor) (40–42). This dual-labeled TDP-43 was immobilized onto PEG-biotinylated glass using an N-terminally encoded HaloTag, isolating individual molecules across the glass surface which can be illuminated via TIRF to observe the conformational behavior of the TDP-43 LCD on these molecules (Figure 3A, B). Additionally we used an alternating laser excitation (ALEX) scheme, where two lasers alternately excite the donor and acceptor dyes, to assess dye stoichiometry and one-step photobleaching in order to select molecules dually labeled with one Cy3 dye and one ATTO647 dye (Figure 3C) (40, 62). FRET efficiency informs on the conformation of the LCD where high FRET occurs when the fluorophores are in close proximity, corresponding to the collapse of the LCD while low FRET occurs when the fluorophores are distant, corresponding to extension of the LCD (Figure 3D). Under incubation in 0.3 mg/mL BSA control, we observed a broad range of FRET signals, with a slight peak at ∼0.8, suggesting high heterogeneity in the distribution of LCD conformations (Figure 3E), in agreement with its disordered nature and tendency to aggregate (63, 64). We also introduced the A315T mutation into the genetic code expanded TDP-43 for single-molecule analysis. Unlike the WT LCD, we observed a strong high FRET peak (∼0.9) for the A315T mutant LCD when incubated in the control BSA-containing buffer (Figure 3F), indicating that the A315T mutation induces more collapsed conformations than WT LCD.

**Figure 3.**
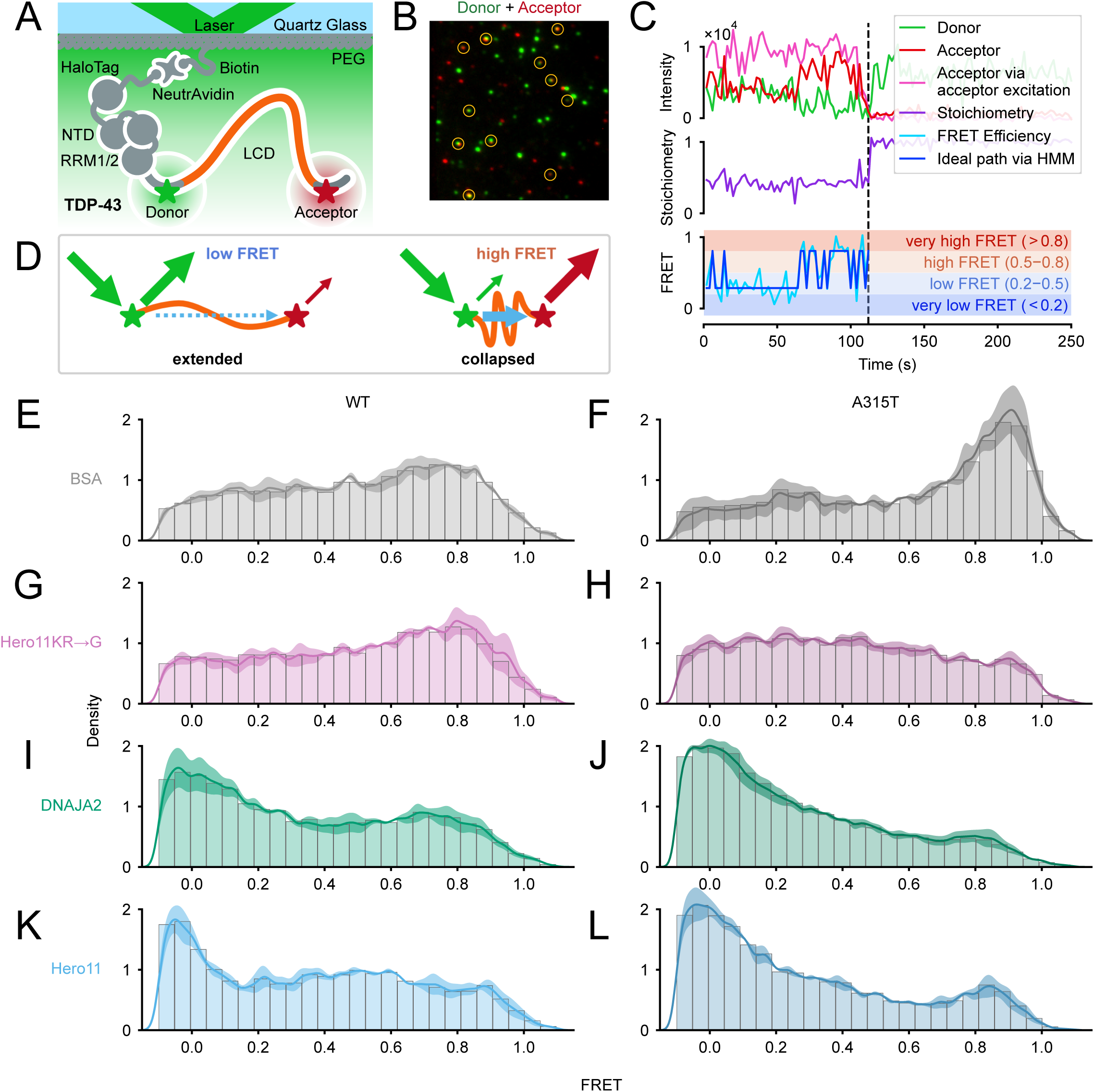
Hero11 and DNAJA2 promote extended conformations of the TDP-43 LCD. (A) Schematic of TIRF-smFRET observation of the LCD (orange) on full-length TDP-43. HaloTag-TDP43 was labeled with Cy3 dye (Donor, green) and ATTO 647N dye (Acceptor, red) around the LCD and immobilized via NeutrAvidin to biotin-PEG on a quartz glass surface. Note that donor and acceptor dyes are randomly incorporated at the two sites, but only molecules dually labeled with one donor and one acceptor were selected for analysis (see Methods), of which the case where the donor is before and acceptor after the LCD is represented here. (B) Representative smFRET microscope image (from DNAJA2 incubation) of molecules tethered to the glass surface. Donor (green) and acceptor (red) emission signals are overlaid here, where candidate single molecules are highlighted in yellow circles. (C) Example FRET trace selected for analysis (from Hero11 incubation). The top plot shows the corrected fluorescence intensities of the Cy3 donor (green) and ATTO647N acceptor (red) signals upon excitation of the donor dye. The fluorescence intensity of the ATTO647N acceptor signal under excitation of the acceptor dye (from alternating-laser excitation) is shown in magenta. The middle plot shows the dye stoichiometry (purple), where roughly constant signals at ∼0.4 are attributed to single molecules dually labeled with one donor and acceptor dye. The bottom plot shows the FRET efficiency (cyan), with the ideal path from a hidden Markov model (blue) overlaid. The shaded background in this plot corresponds to the binning used in the dynamics analyzes in Figure 4. The vertical dotted line shows when the first single step photobleaching event occurred. In this case, the acceptor dye bleaches, resulting in complete loss of acceptor signal in both donor and acceptor excitations, recovery of donor signal in absence of FRET, and loss of ∼0.4 dye stoichiometry. (D) Schematic of how FRET signals provide information on LCD conformation. Upon excitation (green downward arrow), the donor dye Cy3 (green) may fluoresce (green upward arrows) or transfer resonance energy to the acceptor dye, ATTO 647N dye (red) via FRET (blue arrow), exciting the acceptor dye, in turn allowing it to fluoresce (red upward arrows). The transfer efficiency depends on the 6^th^ power of the dye-to-dye distance, resulting in a steep distance dependence for the FRET signal (Supplementary Figure 4A). Left: low FRET occurs when the dyes are far apart, corresponding to extended LCD and is observed as more donor signal compared to acceptor. Right: high FRET occurs when the dyes are near, corresponding to collapsed LCD and is observed as less donor signal compared to acceptor. (E–L) FRET efficiency distributions of TDP-43 LCD WT or TDP-43 LCD A315T incubated with 0.3 mg/mL BSA (E, F), Hero11KR→G (G, H), DNAJA2 (I, J) or Hero11 (K, L), respectively. The density plots represent the mean ± SEM (the solid lines and shaded areas, respectively) of three repeated experiments. Histograms contain all the observations. The total numbers of molecules in each histogram are (E), 150; (G), 160; (I), 178; (K), 168; (F), 101; (H), 255; (J), 357; and (L), 216. Note that values less than 0 and greater than 1 arise from background correction uniformly across molecules of an experiment and from the kernel in the density plot and still correspond to valid observations (see Methods).

### Hero11 and DNAJA2 promote extension of the TDP-43 LCD

Since previous TIRF-smFRET studies of bacterial Hsp40s showed that they stabilize conformationally expanded states of natively folded client proteins such as luciferase and rhodanese (65, 66), we investigated whether eukaryotic DNAJA2 was capable of eliciting this effect on the intrinsically disordered client TDP-43 LCD. In accordance with these studies, incubation of LCD with DNAJA2 resulted in a very low FRET peak at ∼0.0 (Figure 3I), which indicated extension of the LCD conformation. Strikingly, the addition of Hero11 also caused a similar very low FRET peak (Figure 3K). In contrast, Hero11KR→G mutant did not cause apparent changes in the FRET distribution compared with the BSA control (Figure 3G), consistent with its inability to suppress TDP-43 aggregation in cells (Figure 1D). To account for the concentration dependence of these effects, we repeated this experiment at 1/4^th^ lower protein concentration (Supplementary Figure 2). While lowering the concentration of Hero11KR→G (Figure 2A) or DNAJA2 (Figure 2B) from 0.3 mg/mL to 0.075 mg/mL did not notably alter their effects on LCD conformational distributions, lowering the concentration of Hero11 did modestly weaken the ∼0 low FRET peak (Figure 2C). However, the LCD conformational distribution in 0.075 mg/mL Hero11 was nonetheless similar to DNAJA2 at 0.3 mg/mL (and 0.075 mg/mL). Given that Hero11 is around 1/4^th^ the molecular weight of DNAJA2, this suggests that equimolar amounts of Hero11 and DNAJA2 show similar effects on LCD conformation. Nonetheless, these proteins are still expected to be in excess compared to TDP-43, which are isolated as single molecules across the glass surface.

Similarly, consistent with their effectiveness in suppressing the aggregation of the A315T mutant in cells (Figure 1E), both Hero11 and DNAJA2 promoted very low FRET peaks (∼0.0) in the FRET distribution of the A315T LCD (Figure 3J, L). Unlike the WT LCD, where its influence on conformational distribution was indistinguishable from that of the BSA control, incubation with Hero11KR→G resulted in loss of the very high FRET peak (∼0.9), resulting in a flat conformational distribution of the A315T LCD (Figure 3H). This suggests that the KR→G mutant can reduce the population of LCD molecules trapped in very high FRET states, but unlike DNAJA2 and Hero11 WT, did not cause a shift to a low FRET peak. Altogether this suggests that DNAJA2 promotes extended conformations of the aggregation-prone intrinsically disordered WT and A315T TDP-43 LCD, and that Hero11, despite lacking specific tertiary structures such as those of DNAJA2, is capable of eliciting a similar effect in a manner dependent on its net charge.

### Hero11 and DNAJA2 stabilize conformationally dynamic extended states of the LCD

We next investigated the detailed conformational dynamics of TDP-43 LCD in the presence of the aggregation suppressors. To this end, we idealized the dynamic FRET trajectories into discrete FRET states via a hidden Markov model (HMM), allowing for quantitative analysis of conformational dynamics (Figure 3C) (40, 49). To understand how DNAJA2 and Hero11 change transition dynamics, FRET states were classified into 4 bins based on FRET efficiency: very low (<0.2), low (0.2–0.5), high (0.5–0.8), and very high (>0.8). Then, we counted transition events between these FRET bins for all possible combinations, summarizing the frequency of transition events occurring between FRET bins (Figure 4A). While there were some weak transition patterns in BSA (between low and high bins and vice versa) and Hero11KR→G (from very low to high) incubations, there was no clear overall pattern in the effects of these proteins on transition events. In contrast, we observed that most transition events (∼20%) occurred within the very low FRET bin for the DNAJA2 and Hero11 incubations, with little pattern elsewhere. This suggests that both DNAJA2 and Hero11 can maintain the dynamic transitions of the LCD around very low FRET (i.e., the extended state of the LCD as in Figure 3I, K).

**Figure 4.**
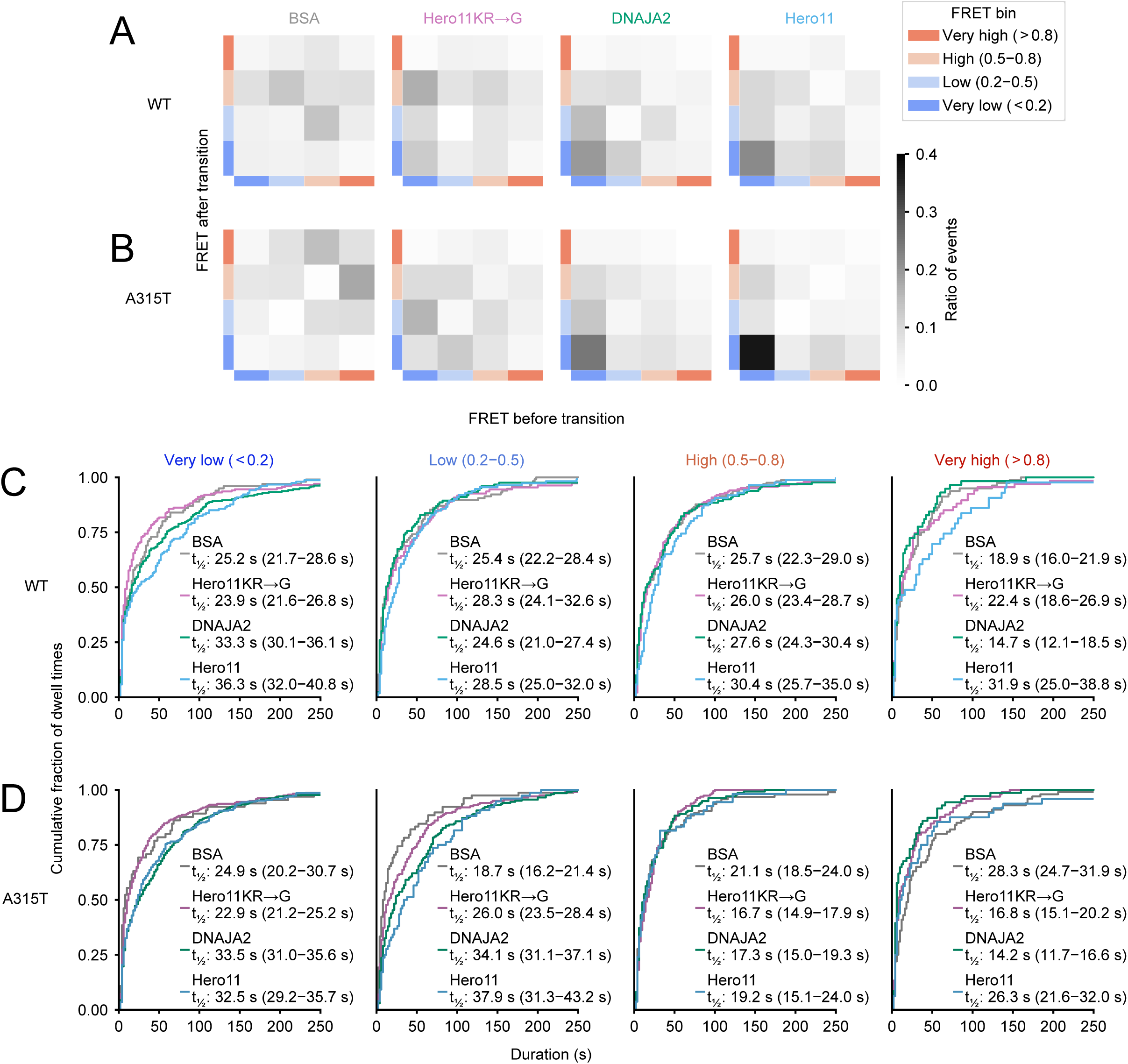
DNAJA2 and Hero11 stabilize the dynamic and extended states of LCD. (A, B) Transition density plots of the WT (A) or A315T (B) LCD, under incubation in the indicated protein, from left to right: BSA, Hero11KR→G, DNAJA2, and Hero11. Idealized FRET traces were obtained from HMM from which FRET states were obtained and binned by FRET efficiency: very low (<0.2), low (0.2–0.5), high (0.5–0.8), and very high (>0.8), as in Figure 3C. Then, the number of transition events between the FRET bins were counted. The ratio of transition events between the FRET state bins is indicated by the shading. (C, D) Dwell times of FRET states as represented by empirical cumulative distribution functions for the WT (C) or A315T (D) LCD grouped by the FRET state bin. From the idealized FRET traces, the dwell times were measured as the duration of each FRET event and grouped by the FRET bin as in Figure 3C, and further grouped by incubation condition as indicated by the color. For each bin, the dwell times under each incubation condition were fitted to single exponential models (see Supplementary Figure 3). From the fitted models, the estimated dwell time half-lives (s), t_1⁄2_, are displayed, and uncertainties reported as 95% credible intervals in the paratheses.

For the A315T mutant (Figure 4B), the transition density histograms in the BSA control display most of the transition events occurring between the high- and very high-FRET bins. Incubation with Hero11KR→G results in evenly distributed FRET state transitions from very low to very high bins, consistent with its flat conformational distribution (Figure 3H). In contrast, for Hero11 or DNAJA2, transition events occurred more frequently within the <0.2 FRET bins, suggesting the dynamic behavior of molecules in the very low FRET peak promoted by these proteins (Figure 3J, L). Taken together, the effects of BSA, Hero11, Hero11KR→G, and DNAJA2 on the transition dynamics of the A315T mutant LCD were generally consistent with their effects on the WT LCD.

We next measured the dwell times of the FRET states (duration spent in a certain FRET state before transitioning away to another state) for the four bins (very low to very high, as shown in Fig. 4A and B), summarized as empirical cumulative distribution functions (Figure 4C, D). Because transitions between the four FRET bins are expected to be governed by single-step processes, we used a Bayesian approach to fit single exponential distribution models to each of the dwell time distributions to quantitatively sample/estimate the kinetic rate constant (reported as the dwell time half-life, t_½_; see Supplementary Figure 3 for fitting accuracy).

For the WT LCD (Figure 4C), Hero11 and DNAJA2 clearly extended the dwell times of the very low FRET states of the LCD compared to BSA and Hero11KR→G (median t_1⁄2_ ∼33–36 s vs ∼24–25 s, for Hero11 and DNAJA vs BSA and Hero11KR→G, respectively). Similarly, Hero11 and DNAJA2 also extended the dwell times of the very low FRET states of A315T compared to BSA and Hero11KR→G (Figure 4D; median t_1⁄2_ ∼33 s vs ∼23–25 s, respectively). Additionally, this extension of dwell time also occurred for the low-FRET states of A315T (median t_1⁄2_ ∼34–38 s vs ∼19–26 s for Hero11 and DNAJA2 vs BSA and Hero11KR→G, respectively). This suggests that Hero11 and DNAJA2 stabilize the extended conformations of both the WT and A315T LCD.

Taken together, our detailed dynamics analyses indicate that Hero11 and DNAJA2 both stabilized the dwell times of the lowest FRET states (Figure 4C and D) while biasing for transition events between these lowest FRET states (Figure 4A and B) of the WT and A315T LCDs. For DNAJA2, this effect is consistent with the stabilization of unfolded client proteins by Hsp40s (24, 65, 66). We also found that Hero11 has similar effects despite the lack of the structural features on Hsp40 necessary for interaction with TDP-43 LCD (24, 56). Thus, despite differences in structural features, these results indicate that Hero11 and DNAJA2 have surprisingly similar effects on the dynamics of the TDP-43 LCD.

## Discussion

Overall, our results orchestrally support that conformational modulation of the LCD is a key to the pathological aggregation process of TDP-43. Although it is currently almost impossible to directly observe conformation within aggregates via smFRET, we envision that these compact conformations, which are amenable to extension, may represent precursors to stably aggregated states. In support of this, we estimate the very high FRET ∼0.9 peak corresponds to an ∼3.3 nm mean end-to-end distance for our dye pair, labeled at residues 268 and 420 six amino acids before and after the LCD (Figure 4A), which is roughly compatible with the distances in collapsed LCD conformations observed in amyloid fibril structures of the in vitro-formed full LCD (7KWZ [PDB code]: F276–M414, 1.716 nm) (14). This suggests the A315T mutation induces more compact conformations of LCD at the single-molecule level in addition to its role in stabilizing the amyloid fibril structure through intra- and interstrand interactions (16, 17).

In contrast, we observed that DNAJA2 stabilized extended conformations of the TDP-43 LCD and suppressed its aggregation. This finding is in line with reports that Hsp40s stabilize nonnative unfolded conformations of structured client proteins, leading to native-state folding (24, 65–67), and reports that DNAJA2 binds to the LCD (56). However, the distinction here is that the client protein is an otherwise collapsed LCD. We also found that Hero11 stabilized the extended conformations of the TDP-43 LCD similarly to DNAJA2. In support of this, small but increasing numbers of Hero11 molecules lead to an increased radius of gyration for LCD molecules in molecular dynamics simulations (68). The very low FRET ∼0 peak observed in the presence of Hero11 and DNAJA2 should contain conformations from 6.80 nm (0.1 FRET) to wider than 10.2 nm (0.01 FRET) mean end-to-end distance (∼0.01 FRET), which is comparable or more extended than the ∼7.34 nm mean distance expected for a “native” state of the LCD, estimated by a wormlike chain model (Supplementary Figure 4A, B; see Methods). Together, these support conformational modulation of the LCD as a determinant of TDP-43 aggregation, where extended conformations, promoted by the intervention of DNAJA2 or Hero11, are excluded from progressing further toward aggregation, as expected for “holdase”-type Hsp40 chaperoning (21, 22, 69).

Despite the similar effects on TDP-43 LCD conformation, a key difference between DNAJA2 and Hero11 is that DNAJA2 extensively utilizes its structural features, including multivalent binding at multiple sites across the surfaces of its C-terminal domains across the dimer, in its chaperoning of client proteins (24). In contrast, Hero11 is unlikely to have a solid tertiary structure or bind to TDP-43 LCD in a defined conformation. Nonetheless, it is possible that Hero11 interacts with the TDP-43 LCD via a partial alpha-helical secondary structure on both IDPs, as demonstrated in molecular dynamics simulations (68). Notably, such interactions between Hero11 and the LCD made up a large fraction of the intermolecular contacts in these simulations (68). The transient alpha-helical structure in the LCD is also selectively bound by DNAJA2 (56), which may serve as an anchoring interaction for stabilizing LCD conformation. Alternatively, but not mutually exclusively, this stabilization effect may occur based on the biophysical or compositional properties of the Hero11 chain regardless of structure because exclusion of this alpha-helical structure did not substantially affect the overall results of the simulations (68). As demonstrated in designed random polymers mimicking IDPs (70), the conformational flexibility and composition of Hero11 may maximize stabilizing interactions with the surfaces of client proteins to prevent aggregation. Notably, the highly positively charged property of Hero11 is important for its interaction with the LCD, as the charge composition mutant Hero11KR→G failed to elicit stabilization of the extended conformations nor suppress aggregation. Compositional charge may be a requirement for this effect as other positively charged hero proteins, Hero45 and Hero7 (pI values 8.66 and 10.44, respectively) were also effective suppressors of TDP-43 aggregation, while negatively charged hero proteins, Hero9, Hero20 and Hero13 (pI values 4.13, 4.57 and 5.31, respectively) were poor suppressors (36). Thus, we speculate that the combination of transient interactions and the compositional properties of Hero11 enable it to influence client protein conformation.

Moreover, these biophysical properties of Hero11 may enable it to suppress phase separation of the LCD (and presumably subsequent aggregation) according to molecular dynamics simulations (68). This curiously mirrors how DNAJA2 suppresses LCD phase separation, where cationic residues of DNAJA2 are thought to supply similar repulsive effects (56). However, one key difference is that DNAJA2 is a co-chaperone of Hsp70 (20, 56), allowing for recruitment of TDP-43 towards the greater chaperoning network, whereas such activity is still unknown for Hero11. Nonetheless, overexpression of Hero11 has been demonstrated to bolster the fitness of cultured cells and *Drosophila* (36), presumably by reducing the burden of immediately harmful misfolded proteins on the canonical chaperoning system. Understanding the downstream mechanistic details of hero protein-mediated aggregation suppression will be important not only from a basic science perspective but also for their biotechnological and therapeutic applications.

## Acknowledgments

We thank all members of the Tomari laboratory for critical comments on the manuscript.

