## Supplementary Figure S1-S4 for "DNAJA2 and Hero11 mediate similar conformational extension and aggregation suppression of TDP-43"

### Supplementary Figure 1

A

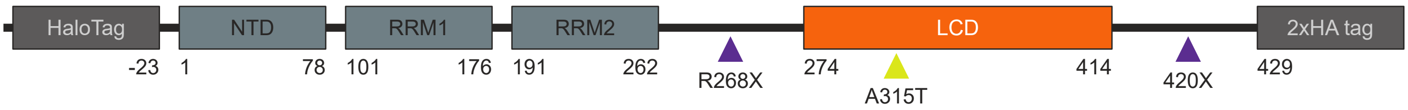


B

```
GMAEIGTGFPFDPHYVEVLGERMHYVDVGPRDGTPVL
FLHGNPTSSYVWRNIIPHVAPTHRCIAPDLIGMGKSD
KPDLGYFFDDHVRFMDAFIEALGLEEVVLVIHDWGSA
LGFHWAKRNPERVKGIAFMEFIRPIPTWDEWPEFARE
TFQAFRTTDVGRKLIIDQNVFIEGTLPMGVVRPLTEV
EMDHYREPFLNPVDREPLWRFPNELPIAGEPANIVAL
VEEYMDWLHQSPVPKLLFWGTPGVLIPPAEAARLAKS
LPNCKAVDIGPGLNLLQEDNPDLIGSEIARWLSTLEI
SGHRYTSLYKKAGSAAELGTLEMSEYIRVTEDENDEP
IEIPSEDDGTVLLSTVTAQFPGACGLRYRNPVSQCMR
GVRLVEGILHAPDAGWGNLVYVVNYPKDNKRKMDETD
ASSAVKVKRAVQKTSDLIVLGLPWKTTEQDLKEYFST
FGEVLMVQVKKDLKTGHSKGFGFVRFTEYETQVKVMS
QRHMIDGRWCDCKLPNSKQSQDEPLRSRKVFVGRCTE
DMTEDELREFFSQYGDVMDVFIPKPFRAFAFVTFADD
QIAQSLCGEDLIIKGISVHISNAEPKHNSNXQLERSG
RFGGNPGGFGNQGGFGNSRGGGAGLGNNQGSNMGGGM
NFGAFSINPAMMAAAQAALQSSWGMMGMLASQQNQSG
PSGNNQNQGNMQREPNQAFGSGNNSYSGSNSGAAIGW
GSASNAGSGSGFNGGFGSSMDSKSSGWGMLEGSEXKL
VDLQSRYPYDVPDYASRYPYDVPDYASR*
```

768 amino acids
~85 kDa

C

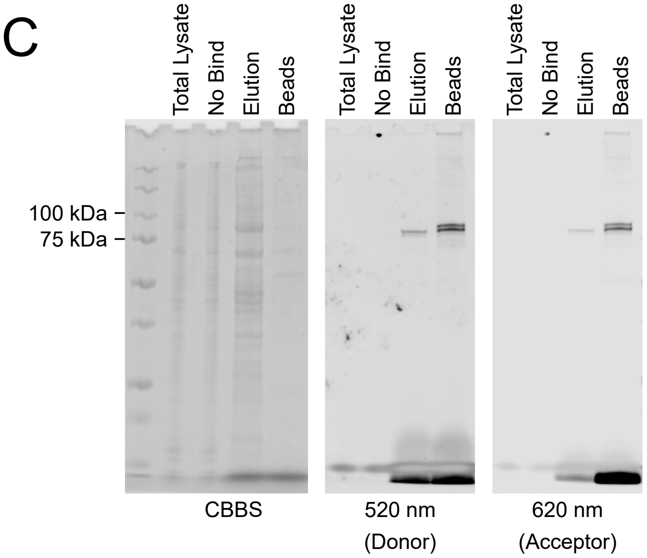

### Supplementary Figure 2

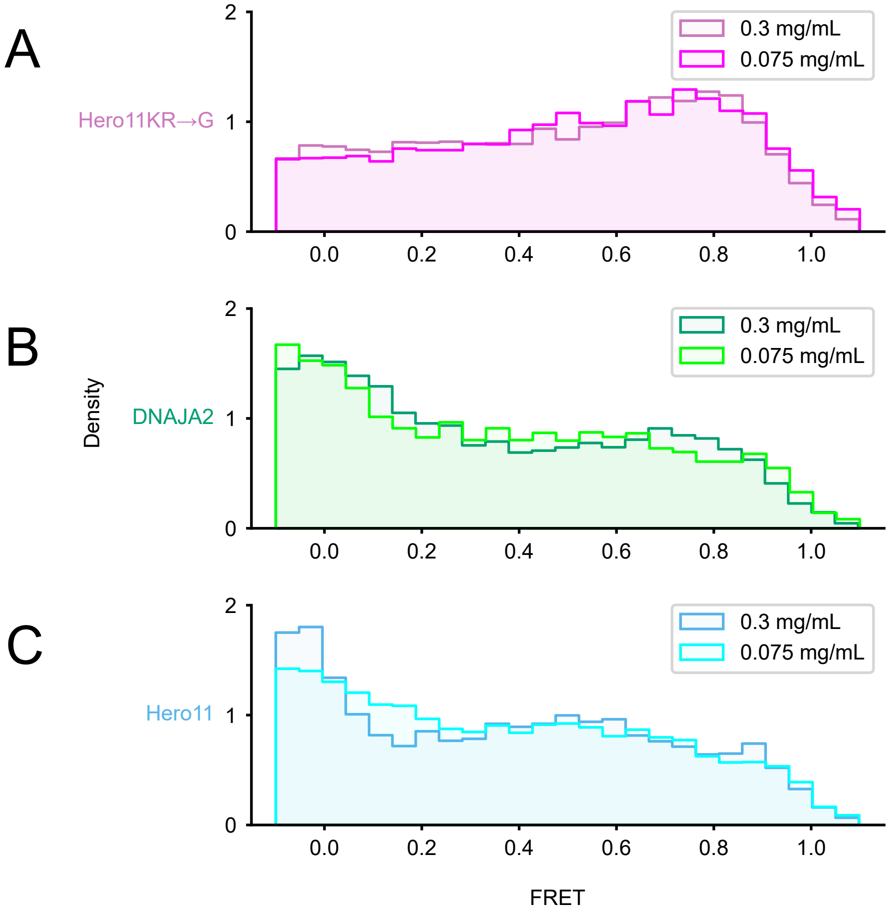

### Supplementary Figure 3

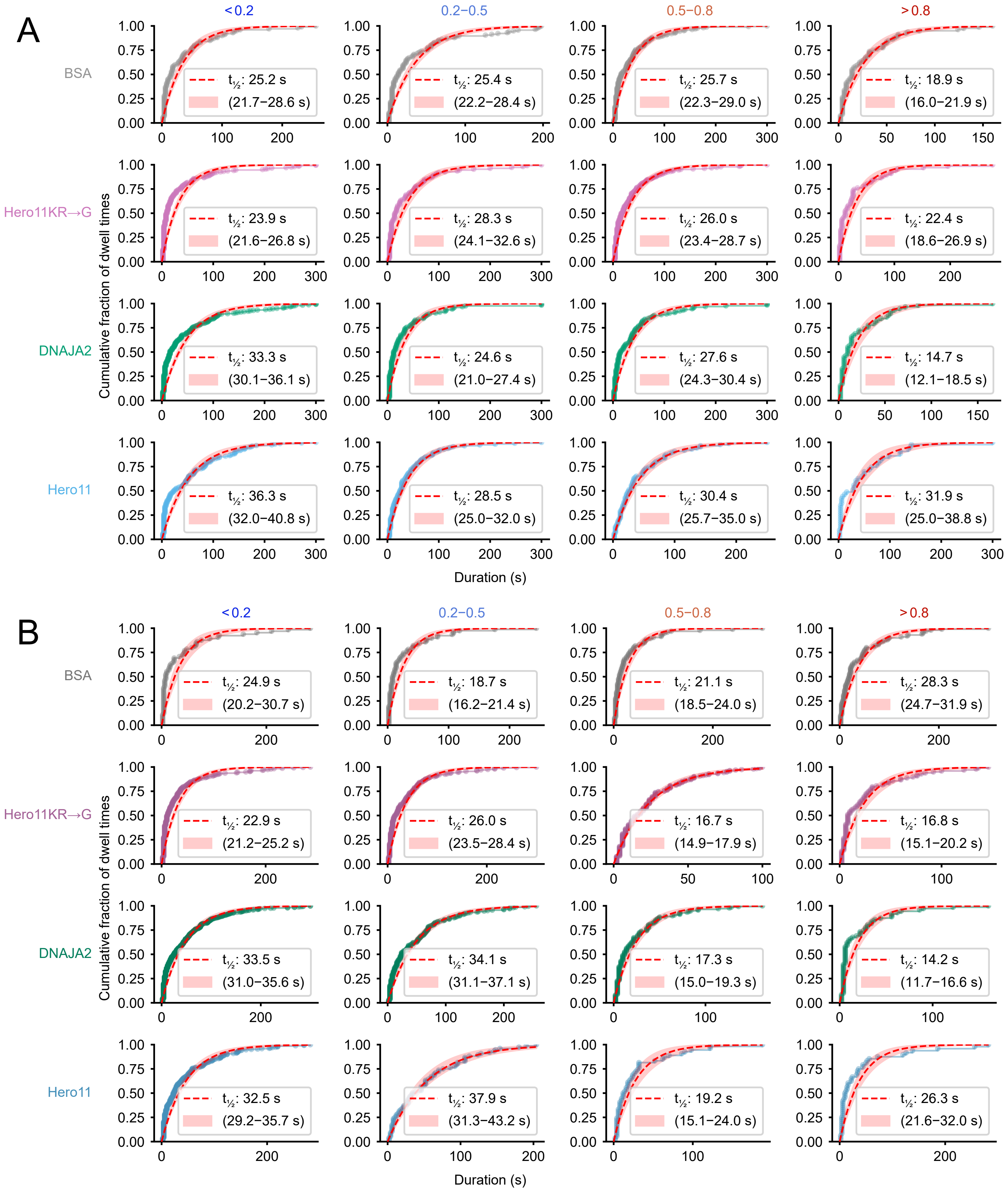

### Supplementary Figure 4

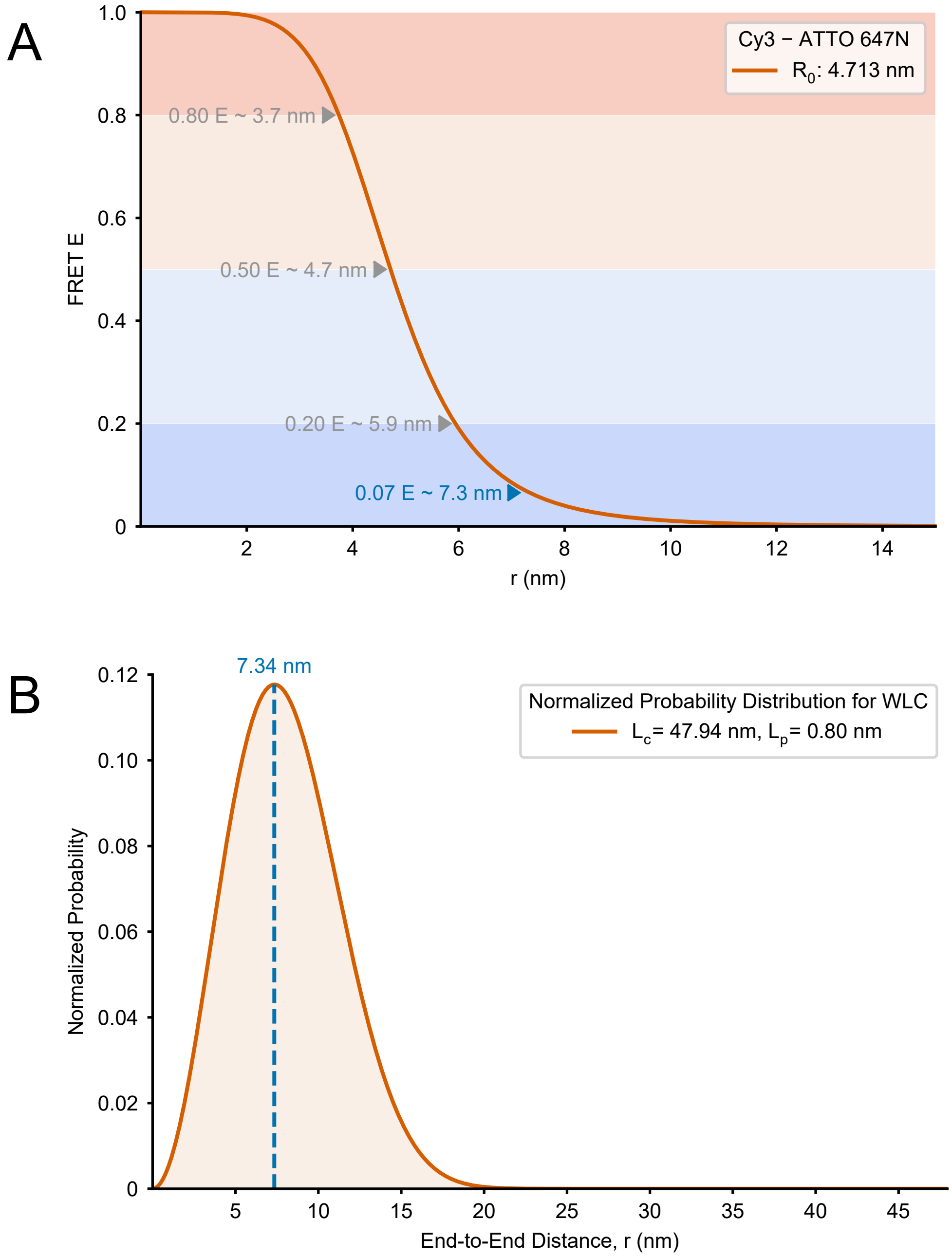
