## Supplementary Figure Legends for "DNAJA2 and Hero11 mediate similar conformational extension and aggregation suppression of TDP-43"

### Supplementary figure text

#### Supplementary Figure 1. Genetic code expansion in HEK293T cells for dye labeling of TDP-43

(A) Schematic of HALO-TDP43 with unnatural amino acids incorporated around LCD after elution via TEV protease cleavage. Boxes indicate TDP-43 regions and tags: the N-terminal domain (NTD), two RNA recognition motifs (RRM1 and RRM2), a low-complexity domain (LCD), a HaloTag, and a 2xHA tag. The last amino acid of HaloTag is the 23^rd^ amino acid from the start of the TDP-43 sequence. Sites of unnatural amino acid (endo BCN-L-lysine) incorporation are indicated by violet triangles, where arginine 268 was replaced with unnatural amino acid (R268X), while the other site was incorporated at residue 420 (420X; the sixth residue after the end of the TDP-43 sequence, residue 414) in the linker before the 2xHA tag. The yellow triangle marks the site of the A315T mutation.

(B) The amino acid sequence of (A). The highlighted color corresponds to the regions in (A). X represents unnatural amino acid, and the A highlighted in yellow is alanine 315, which is mutated to threonine in the A315T mutant. The N-terminal glycine residue is left over from the TEV protease cleavage, and the asterisk marks the end of the sequence.

(C) Dye-labeling of HALO-TDP43. HEK293T cells expressing FLAG-tev-HALO-TDP43 incorporated with unnatural amino acids were lysed and incubated on anti-FLAG Dynabeads. The proteins were dye-labeled on beads and eluted via cleavage with TEV protease in naïve HEK293T cell lysate (note the multiple bands in the elution fraction of the CBB stain). Copper-free click chemistry-based dye labeling of endo BCN-L-lysine with tetrazine-Cy3 (Donor) and tetrazine-ATTO-647N (Acceptor) was confirmed by fluorescence imaging in gel via excitation at 520 nm and 620 nm, respectively, before staining with Coomassie Brilliant Blue (CBB). The two bands in the Elution fraction correspond to HALO-TDP43 with only one or two dyes incorporated (lower and upper bands, respectively). The Beads fraction indicates proteins that did not elute from the Dynabeads, and higher molecular weights correspond to dye-labeled but uncleaved FLAG-tev-HALO-TDP43.

#### Supplementary Figure 2. Effects on LCD conformation remain consistent with lower concentrations of DNAJA2, Hero11, or Hero11KR→G

(A – C) FRET efficiency histograms of the TDP-43 LCD WT incubated in 0.075 mg/mL Hero11KR→G (A), DNAJA2 (B), or Hero11 (C), compared against 0.3 mg/mL. BSA was supplemented for the same total protein concentration of 0.3 mg/mL to accommodate comparison between the different concentrations. Each histogram contains all observations for a single experiment and the total number of molecules in each histogram are (A), 68; (B), 101; (C) 96. Histograms from Figure 3G, I and K, corresponding to 0.3 mg/mL of the respective protein are replotted here for comparison.

#### Supplementary Figure 3. Fitting of exponential distribution models to dwell time distributions

(A, B) Model fits for the empirical cumulative distribution functions for the WT (A) and A315T (B) LCDs in Figure 4C and D. The columns are FRET state bins, and the rows are the protein incubation. Each dot represents an observed dwell time, to which exponential distribution models were fitted to obtain median and 95% credible interval estimates of the rate parameter, which are plotted as the red dashed lines and shaded area.

#### Supplementary Figure 4. Estimation of theoretical LCD distances for the Cy3 – ATTO 647N dye pair

(A) The theoretical relationship between FRET efficiency and distance for the Cy3 – ATTO 647N dye pair with a Förster distance, $R_{0}$, of 4.713 nm. Very low FRET signals (below 0.2) are expected to correspond to distances greater than ~6 nm.

(B) The theoretical probability distribution of end-to-end distances, $P(r)$, for a 141 amino acid-long chain following wormlike chain behavior, using polymer contour length of 47.94 nm and persistence length of 0.80 nm. Peak probability density is centered around 7.34 nm.
